# Nucleus accumbens dopamine release reflects the selective nature of pair bonds

**DOI:** 10.1101/2022.11.10.516053

**Authors:** Anne F. Pierce, David S.W. Protter, Gabriel D. Chapel, Ryan T. Cameron, Zoe R. Donaldson

## Abstract

In monogamous species, prosocial behaviors directed towards partners are dramatically different from those directed towards unknown individuals and potential threats. Dopamine release in the nucleus accumbens plays an important role in guiding social behavior, however, its role in real time social decision making in a monogamous species remains largely unknown. We used monogamous prairie voles to investigate how dopamine release differs in voles when seeking and interacting with a pair bonded partner or a novel vole. Employing the sub-second temporal resolution of the fluorescent biosensor, GRAB_DA_, during a social operant task, we found that partner seeking, anticipation, and interaction resulted in more dopamine release than the same events directed towards a novel vole. Furthermore, partner-elicited dopamine release decreases after prolonged partner separation. Thus, differences in partner- and novel-elicited dopamine release reflect the selective nature of pair bonds and may drive the social behaviors that reinforce and cement bonds over time, eroding after partner loss to facilitate new bond formation.

## Introduction

Optimally navigating social interactions is critical for survival and reproduction, and altered social interaction and cognition contribute substantially to the disease burden of several neurodevelopmental and psychiatric illnesses. Across species, dopamine plays an important role in navigating social relationships. Dopamine is released in the nucleus accumbens during social interaction, and manipulations that increase or decrease dopaminergic system activity within this region promote or impair social interactions, respectively (*1*–*3*). Yet studies to date have examined realtime dopamine dynamics exclusively in laboratory mice (*1, 2*) and rats (*4*), species that exhibit affiliation but do not form selective attachments. Thus, a central question remains of how differences in dopaminergic signaling may contribute to the selective pair bonds that are a cornerstone of the human experience.

Prairie voles are monogamous rodents that form life-long mating-based pair bonds characterized by a selective preference to affiliate with a single partner, as well as aggression towards novel voles of either sex (*5, 6*). Both of these features of pair bonds have been shown to depend on dopaminergic signaling (*7*–*11*). The rewarding nature of pair bonds is reflected in the partner preference test, the primary task used to query prairie vole bond formation (*12*). In this test, a pair bonded vole will choose to spend most of their time with their mating partner compared with a novel opposite sex vole or time spent alone (*12, 13*). Similarly, voles will lever-press to gain transient access to their partner in an operant task (*14*–*16*). However, it remains unknown how real-time dopamine release dynamics contribute to the selective nature of pair bonds and potentially drive partner-directed versus novel-vole-directed behaviors.

To test the hypothesis that highly rewarding partner interaction is accompanied by enhanced dopamine release, we used a novel operant paradigm in which female prairie voles press a lever to gain access to their partner or to a novel opposite sex vole. This task is uniquely advantageous in that it separates the appetitive aspect of social reward, reflected in lever pressing and cues predicting social access, from the consummatory aspect of physical social interaction. Furthermore, this task elicits comparable degrees of lever-pressing for both partner and novel voles (*15*). Using fiber photometric recordings of the fluorescent dopamine sensor, GRAB_DA_, we found that anticipation of partner reunion, as well as partner interaction, results in enhanced nucleus accumbens dopamine release compared with the same events directed towards a novel opposite-sex vole. Further, we found that these differences in dopamine release are labile and can be eroded by long periods of partner separation. Together this suggests that enhanced dopamine release may contribute to pair bond maintenance, and its subsequent erosion upon partner separation may help enable formation of new bonds upon the loss of a partner. These findings are consistent with a reward valuation role for dopamine during social interactions.

## Results

### Dopamine dynamics reflect social operant learning

We performed fiber photometry to measure GRAB_DA_-mediated fluorescence as a proxy for extracellular dopamine levels in the nucleus accumbens of voles engaged in operant responding and reward consumption (Fig 1A - C) (*17*). We used a previously established social operant task in voles (*15*). Voles learned to initially associate rewards with lever pressing through food delivery (Fig S1A - D) before being presented with separate levers that provided transient access to a tethered partner and novel animal (Fig 1D - F).

**Figure 1.**
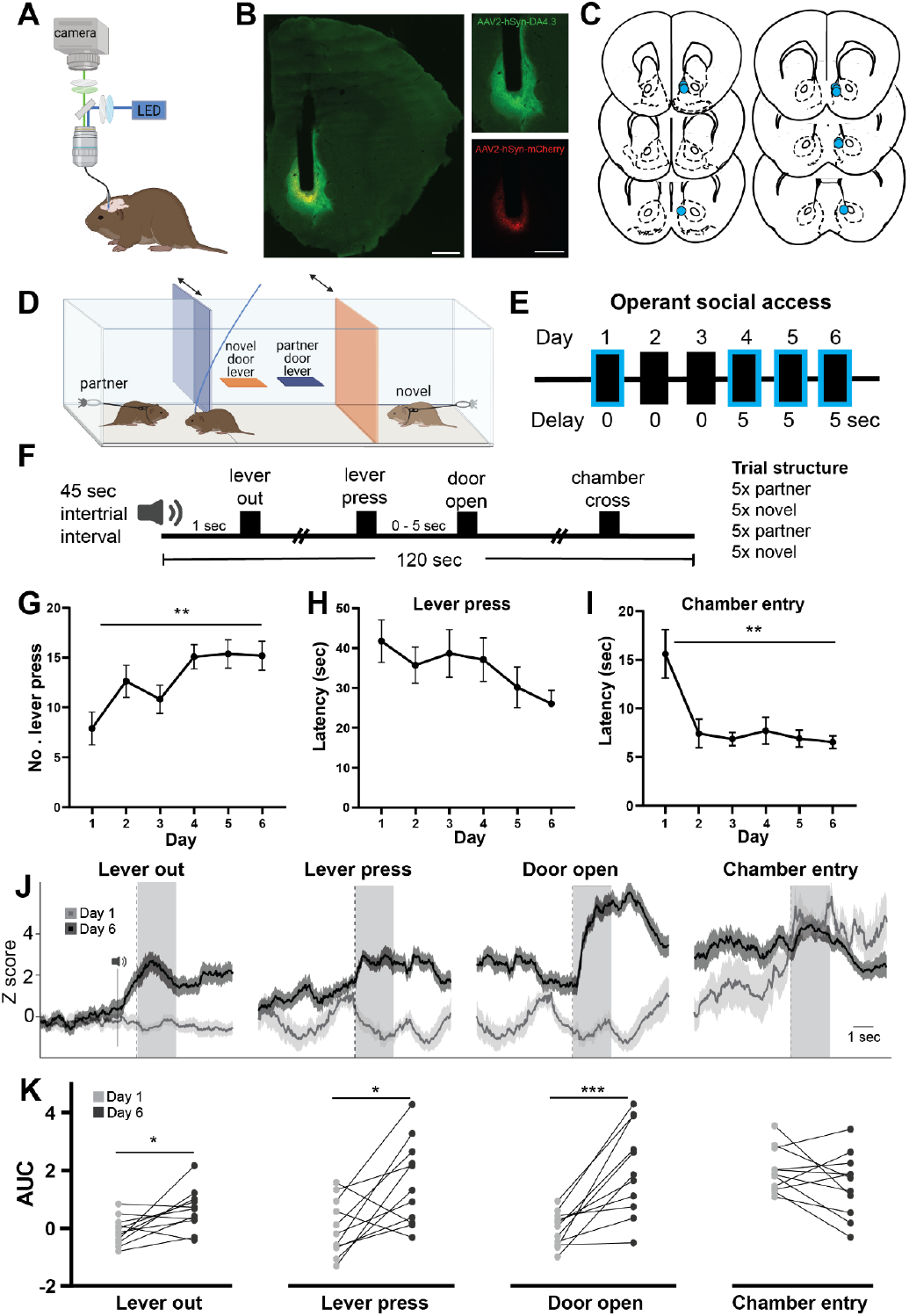
Dopamine dynamics reflect social operant learning. A) Schematic of fiber photometry in a prairie vole. B) GRAB_DA_ and mCherry expression in the nucleus accumbens shell with ferrule track. Scale bar = 500 μm. C) Coronal atlas sections with locations of injection sites and ferrules represented by blue dots. D) Diagram illustrating social operant chamber. E) Experimental timeline of operant social access. Blue outline indicates fiber photometry recording of dopamine levels. Delay indicates the temporal delay (0 or 5 seconds) from lever press to chamber opening. F) Timing of a single trial: A tone is played and either the partner or the novel lever is extended. Subjects have up to 120 s to press the extended lever to open the corresponding social chamber. Subjects underwent 20 trials per day, altering partner and novel access every 5 trials. G) The average number of lever presses for social access increases across training days (one-way repeated-measures ANOVA p = 0.0047) and stabilizes across the last three days of the task. H) Latency to lever press for social access. There is a non-significant decrease in latency to lever press as voles learn the task (mixed model ANOVA p = 0.1783). I) Latency to cross into the social chamber decreases as a function of training day (mixed model ANOVA p = 0.0018). J) Representative GRAB_DA_ fluorescence on the first and last day of operant social access in response to lever extension (lever out), lever press, social chamber opening (door open), and while crossing into the social chamber as detected by an infrared beam break (chamber entry). The onset of each event is indicated by a dashed line. Fluorescence is Z-scored relative to a baseline. K) Area under the curve (AUC) for 2 seconds post-event (shaded regions of J) comparing the first and last training days. (Paired t-tests: Lever out: p = 0.0217; Lever press: p = 0.0146; Chamber open: p = 0.0008; Chamber cross: p = 0.1711). * = p < 0.05; ** = p < 0.01; *** = p < 0.001; n = 11.

In order to determine how dopamine dynamics change as a function of learning in a social operant task, we compared dopamine release on Day 1 and Day 6 of social operant access (Fig 1). Task learning was reflected in increased lever pressing across days (Fig 1G; one-way RM-ANOVA F_(5, 10)_ = 6.139, p = 0.0047), a non-significant decrease in latency to press the lever (Fig 1H; mixed model ANOVA F_(5, 10)_ = 1.9, p = 0.178), and decreasing latencies to enter the social chamber after the door opened (Fig 1I; mixed model ANOVA F_(5, 10)_ = 8.217, p = 0.0018). We found that dopamine release increased for operant events associated with social access on Day 6 relative to Day 1, consistent with prior reports indicating that dopamine release increases as a function of reward anticipation (*2, 18*). Specifically, we observed a significant increase in dopamine following lever extension, lever press, and door opening on Day 6 compared to Day 1 of social operant access (Fig 1J,K; paired t-tests: lever out t_(10)_ = 2.716, p = 0.0217; lever press t_(10)_ = 2.948 p = 0.0146; chamber open t_(10)_ = 4.696, p = 0.0008). We did not observe any differences in dopamine levels associated with chamber entry (Fig 1 J, K, paired t-test; chamber entry t_(10)_ = 1.474, p = 0.1711), which is likely a highly salient event associated with social reward delivery.

### Seeking and interacting with a pair bonded partner elicits enhanced dopamine release

We next asked whether dopamine dynamics distinguished between partner- and novel-associated operant events. We compared intra-animal dopamine release in response to lever extension, lever press, door opening, and chamber entry during interleaved trials in which lever pressing resulted in partner or novel access (Fig 2A). In accordance with prior reports (*15*), voles will press equally and with similar latencies for access to a partner or novel vole in a forced choice context (e.g. when only one lever is available in any given trial) (Fig 2C; 2-way RM-ANOVA: Main effect of vole (partner vs novel): F_(1,10)_ = 1.592, p = 0.2357; Main effect of days F_(5, 50)_ = 6.139, p = 0.0002. Fig 2D; Mixed model ANOVA: Main effect of vole (partner-vs novel): F_(1,10)_ = 1.683, p = 0.159; Main effect of days F_(5, 50)_ = 0.8228, p = 0.2185). Similarly, for partner and novel trials, subjects have similar latencies to enter the social chamber (Fig 2E; Mixed model ANOVA: Main effect of vole (partner vs novel): F_(1,10)_ = 0.5406, p = 0.4791; Main effect of days F_(5, 50)_ = 10.4, p < 0.0001). Across the last three days of social operant access, when operant behaviors were consistent, we observed greater dopamine release for partner lever pressing and door opening compared to the same events in novel trials (Fig 2F and 2G; paired t-tests lever press t_(10)_ = 2.791, p = 0.0191; chamber open t_(10)_ = 2.307, p = 0.0438). Differences in dopamine release for partner and novel trials were not evident after lever extension or in the two seconds after chamber entry (Fig 2F and 2G; paired t-tests lever out t_(10)_ = 1.382, p = 0.1972; chamber entry (0-2s) t_(10)_ = 1.515, p = 0.1606).

**Figure 2.**
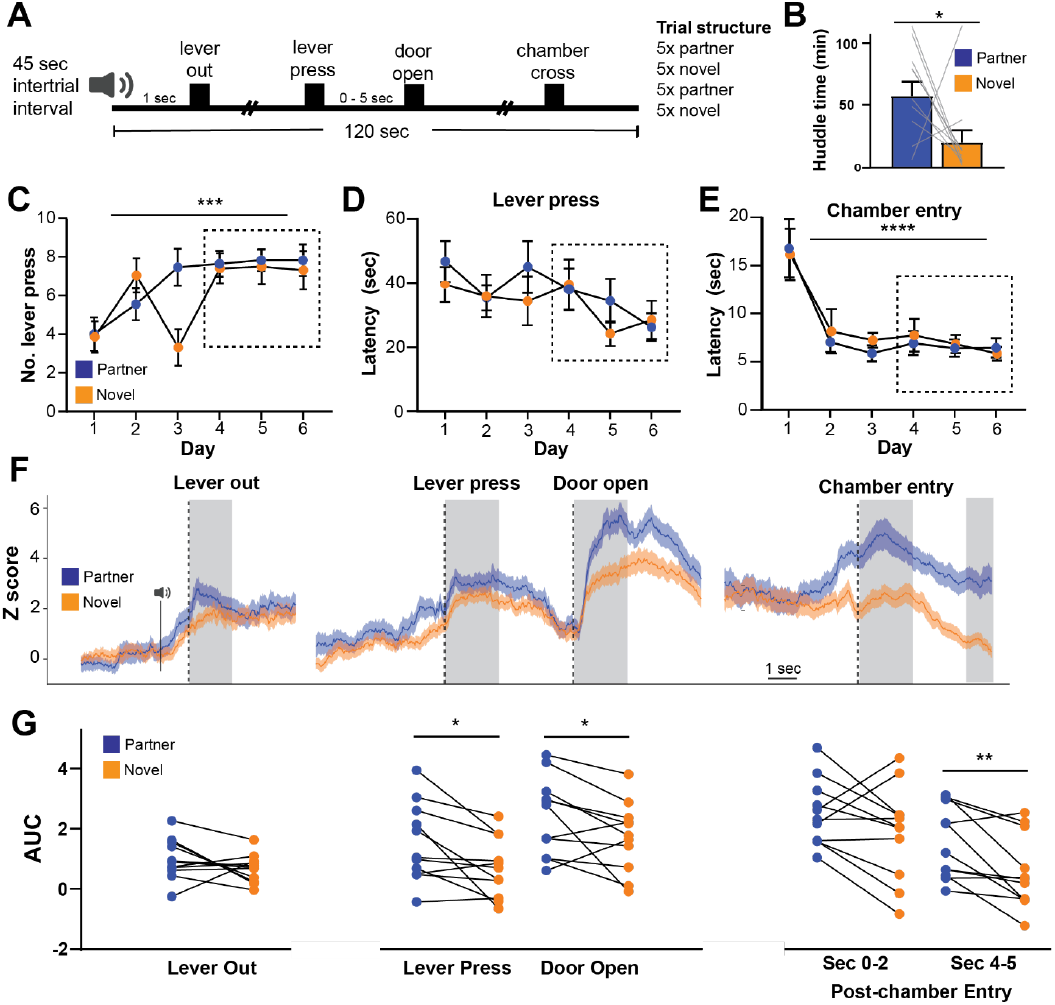
Anticipation of partner access elicits enhanced dopamine release. A) Timing of a single trial. B) Subjects spent more time huddling with their partner compared to the novel in the partner preference test performed after social operant access (one-way t-test relative to no preference (50%), p = 0.0160). C) Number of lever presses for partner and novel access increases across training days. Pressing behavior stabilizes across the last 3 days of operant access and does not differ between partner and novel presses (two-way repeated-measures ANOVA for day p = 0.0002 and partner versus novel p = 0.2357). D) There is a non-significant decrease in latency to lever press across training days and does not differ for partner and novel (two-way repeated-measures ANOVA for day p =0.2185 and partner versus novel p = 0.156). E) Latency to enter the partner or novel chamber decreases as a function of training day but does not differ for partner and novel, and stabilizes by day 2 (two-way repeated-measures ANOVA for day p = <0.0001 and partner versus novel p = 0.4791. F) Representative GRAB_DA_ fluorescence in response to social operant events from days 4 - 6 when operant behavior is consistent. Z-scored fluorescence calculated as in Fig 1J. K) Area under the curve (AUC) for 2 seconds after the event (shaded regions in F) (Paired t-tests: Lever out: p = 0.1972 ; Lever press: p = 0.0191 ; Chamber open: p = 0.0438 ; Chamber cross sec 0-2: p = 0.1606; Chamber cross sec 4-5: p = 0.0086) (* = p < 0.05; ** = p < 0.01; *** = p < 0.001; **** = p < 0.0001) n = 11.

However, when we examined dopamine levels four seconds after social chamber entry, we found there were increased levels of dopamine during partner trials (Fig 2G; paired t-tests chamber entry (4-5s) t_(10)_ = 3.26, p = 0.0086), which prompted us to ask whether behavior and/or dopamine levels differ during social interaction with a partner or novel vole (Fig 3). In accordance with prior reports, voles are more affiliative towards their pair bonded partner, displaying more direct contact, sniffing, and huddling (Fig 3 A,B,E,F; Duration body sniff: paired t-test t_(5)_ = 3.549, p = 0.0164; cumulative body sniff: log-rank (Mantel-Cox) χ2 = 17.67, p = < 0.0001. Duration huddle: unpaired t-test t_(7)_ = 2.423, p = 0.0459; cumulative huddle: log-rank (Mantel-Cox) χ2 = 16.25, p = < 0.0001) (*6*). In contrast, they show greater levels of non-contact investigation towards novel voles (Fig 3 I,J; Duration non-contact investigation: paired t-test t_(10)_ = 2.795, p = 0.019; cumulative body sniff: log-rank (Mantel-Cox) χ2 = 6.565, p = < 0.0001.), potentially serving as an assessment of the conspecific or context. Similar to dopamine release during operant responding to gain social access, we observed greater dopamine release for partner sniffing and huddling compared to the same behaviors directed towards a novel vole (Fig 3 C,D,G,H; body sniff: paired t-test t_(5)_ = 8.974, p = 0.0003; huddle: unpaired t-test t_(7)_ = 3.268, p = 0.0137). We did not observe differences in dopamine elicited by the partner or a novel vole during non-contact investigation (Fig 3 K,L; paired t-test t_(10)_ = 1.085, p = 0.3033).

**Figure 3.**
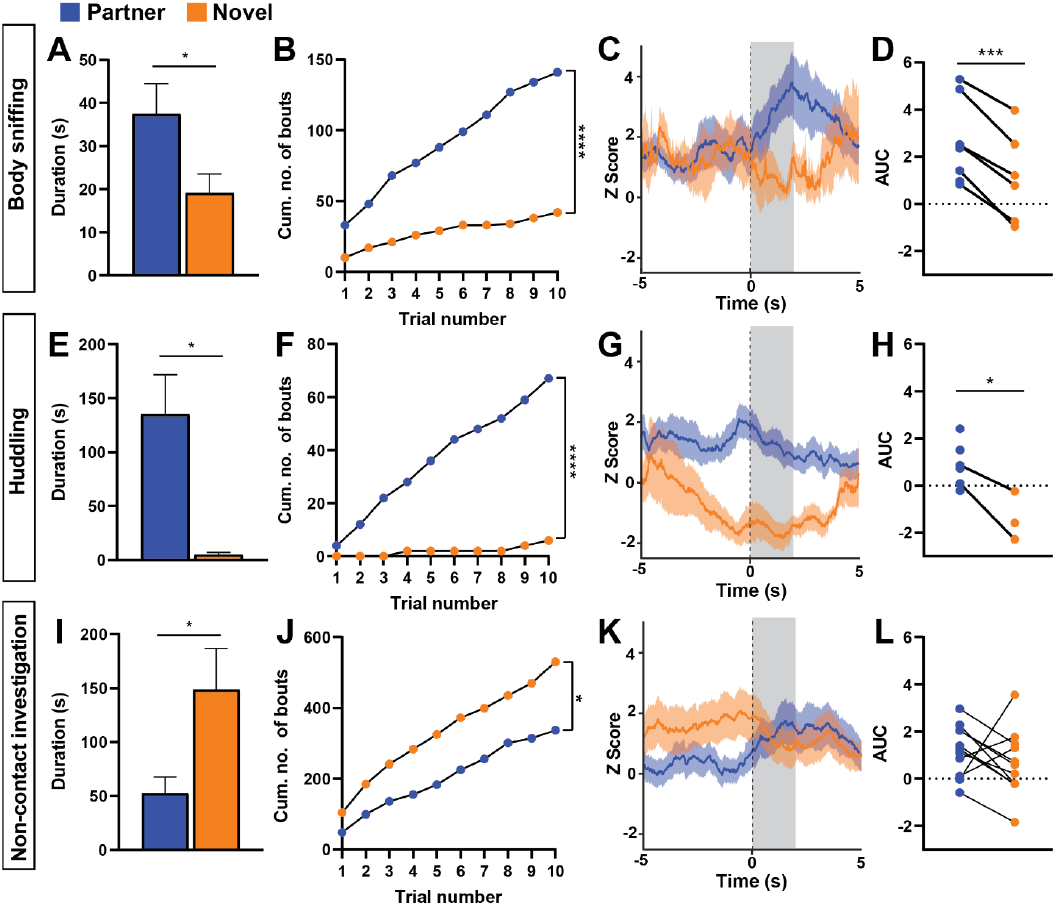
Differences in partner and novel-directed interaction reflect differences in dopamine release. Behaviors displayed toward partners and novels differed following social chamber entry. A - D) Body sniffing: Subjects exhibit greater duration (A) and number of cumulative bouts (B) of contact sniffing towards their partner compared to novel voles (duration: paired t-test p = 0.0164; cumulative bouts: Log-rank (Mantel-Cox) test p = <0.0001). This is associated with greater dopamine release during partner sniffing compared to novel sniffing. Representative trace shown in C with area under the curve calculated for the shaded region (2 sec after behavior initiation) shown in D (paired t-test: p = 0.0003). Z-score calculated relative to baseline as in Fig 2. (E - H) Huddling: Subjects spent more time huddled with their partner than the novel vole after chamber entry (E,F); (duration: unpaired t-test p = 0.0459; cumulative bouts: Log-rank (Mantel-Cox) test p = <0.0001). There was greater dopamine release during partner huddling compared to novel huddling. Trace (G) and AUC (H) unpaired t-test: p = 0.0137. I-L) Non-contact investigation: Subjects exhibited increased duration and number of cumulative bouts of non-contact investigation towards the novel vole (A, B): (duration: paired t-test p = 0.0119; cumulative bouts: Log-rank (Mantel-Cox) test p = <0.0104). There were no differences in dopamine release comparing partner and novel. Trace (K) and AUC (L) paired t-test: p = 0.3033.) Not all animals engaged in all behaviors, especially huddling, as reflected in the reduced number of dots and lack of connecting lines in some instances where an animal huddled only with the partner or with the novel vole.

To further explore the unique features of pair bonding relative to other types of natural reward, we compared dopamine release during social operant and food operant access (Fig S1). Anticipation of food delivery (lever press and pellet dispense) and consumption of the food pellet resulted in significantly less dopamine release than lever pressing, door opening, and chamber opening, respectively, during partner operant trials. However, dopamine release during lever pressing for food and pellet dispensing were not significantly different than lever pressing and chamber opening for novel access (Fig S1F; lever out: one-way RM-ANOVA F_(2, 10)_ = 1.224, p = 0.3122; lever press: one-way RM-ANOVA F_(2, 10)_ = 6.338, p = 0.0079 with Tukey’s post-hoc partner vs food p = 0.0135; novel vs food p = 0.8184; partner vs novel p = 0.0461; chamber open: one-way RM-ANOVA F_(2, 10)_ = 4.75, p = 0.0304 with Tukey’s post-hoc partner-vs food p = 0.0174; novel-vs food p = 0.6368; partner-vs novel p = 0.1009; chamber entry: one-way RM-ANOVA F_(2, 10)_ = 6.005, p = 0.0091 with Tukey’s post-hoc partner-vs food p = 0.0166; novel-vs food p = 0.1689; partner-vs novel p = 0.3249). Thus, partner access elicits greater dopamine than at least two other motivating events -access to a novel vole and access to food.

### Prolonged partner separation erodes enhanced partner-associated dopamine release

Finally, as prior work has shown that voles can form a new bond after 4 weeks of partner separation (*19*), which serves as a proxy for successfully adapting to partner loss, we asked how prolonged partner separation affects partner-associated dopamine release. Prior to separation we performed a partner preference test to confirm a greater preference to huddle with the partner compared to a novel vole. (Fig 2B; one sample t-test relative to 50%: t_(10)_ = 2.895 p = 0.016). Then we separated pairs for four weeks followed by a probe test consisting of one day of social operant access (20 trials) (Fig 4A). When we compared pre- and post-separation dopamine release, we observed a consistent intra-animal decrease in operant-associated dopamine release upon lever presentation, door opening, and chamber entry for partner trials but no change for novel trials (Fig 4B-E; partner paired t-tests lever out: t_(10)_ = 2.88, p = 0.0164; lever press t_(9)_ = 0.04279, p = 0.9668; chamber open t_(9)_ = 2.403, p = 0.0397; chamber entry t_(9)_ = 2.705, p = 0.0242). This decrease in partner-associated dopamine release resulted in the erosion of the differences in dopamine release during partner and novel trials that were evident while animals remained paired (Fig 4F; paired t-tests lever out: t_(10)_ = 0.0379, p = 0.9705; lever press t_(7)_ = 1.502, p = 0.1769; chamber open t_(7)_ = 0.639, p = 0.5432; chamber entry t_(7)_ = 2.039, p = 0.0808). The lack of change in response pre- and post-separation during novel trials indicates that partner-associated decreases in dopamine are not the product of technical considerations, such as a reduction in GRAB_DA_ fluorescence(Fig 4B-E; novel paired t-tests lever out: t_(10)_ = 0.717, p = 0.4898; lever press t_(7)_ = 0.9486, p = 0.3744; chamber open t_(7)_ = 0.5486, p = 0.6003; chamber entry t_(7)_ = 1.479, p = 0.1826). Together, this suggests that partner separation erodes a key pair-bond associated neuromodulatory phenotype. All sample sizes and comprehensive statistical results, including effect size estimates, are reported in **Supplementary Table 1**.

**Figure 4.**
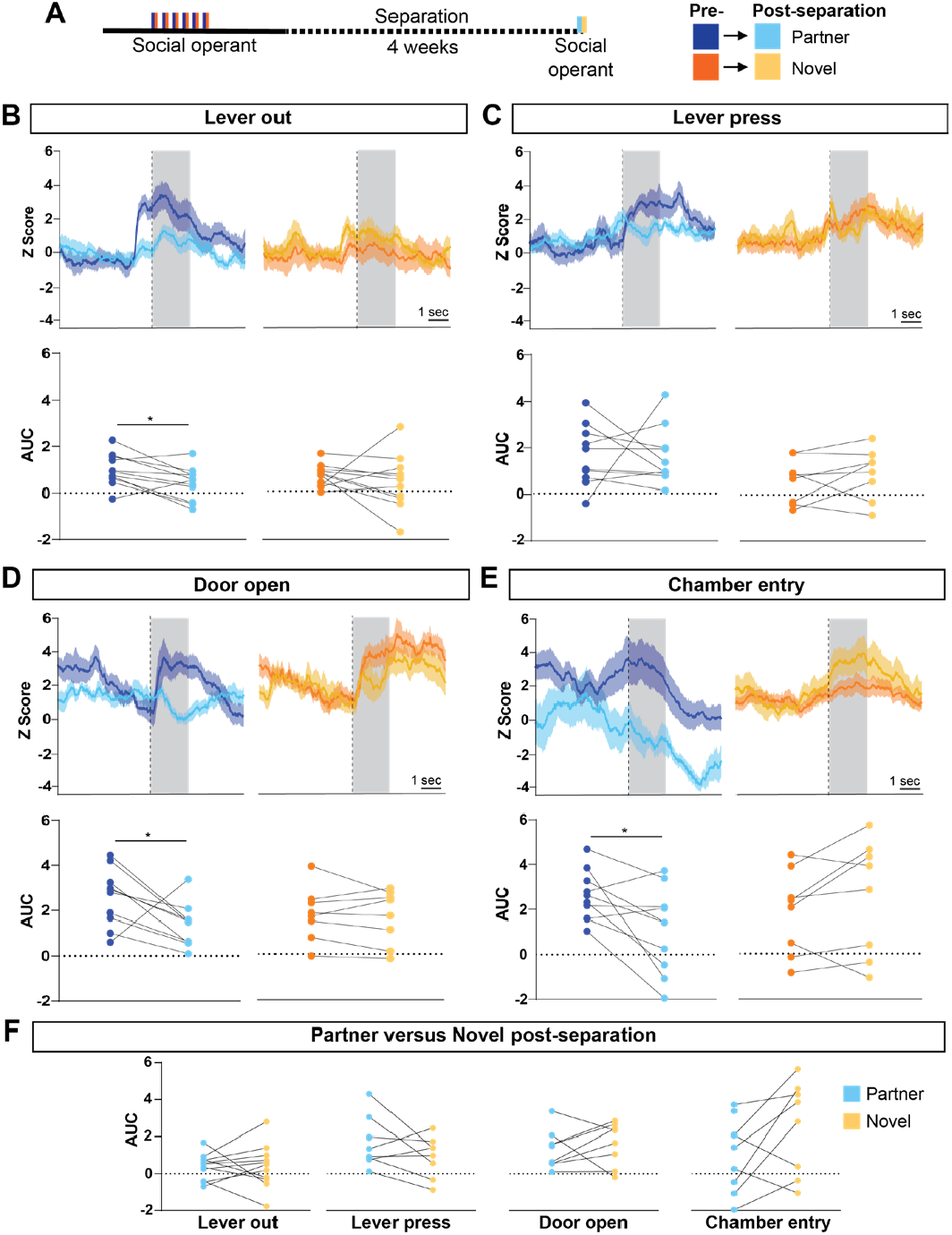
Enhanced partner-associated dopamine levels erode after prolonged partner separation. A) Following social operant access, pairs were separated for 4 weeks, a time period shown to be sufficient to be able to form a new bond that supplants a prior bond. At the end of the 4 week separation, voles underwent a single session of operant social access as in Fig 2A. B - E) Compared to pre-separation, dopamine release was reduced for partner-associated lever out, door opening, and chamber entry while no change was observed for novel-associated events. Top row shows example fluorescent traces with the area under the curve (AUC) shown beneath. Partner pre-post separation paired t-tests: lever out p = 0.0164; lever press p = 0.9668; chamber open = 0.0397; chamber entry = 0.0242. Novel pre-post separation paired t-tests: lever out p = 0.717; lever press p = 0.3744; chamber open = 0.6003; chamber entry = 0.1826. F) The reduction in partner-associated dopamine release is sufficient to erase prior partner-novel differences. AUC for partner versus novel shown for post-separation operant social access as in Fig 2G Partner vs novel post separation paired t-tests: lever out p = 0.9705; lever press p = 0.1769; chamber open = 0.5432; chamber entry = 0.0808. (* = p < 0.05).

## Discussion

Prior pharmacological studies have demonstrated a role for nucleus accumbens dopamine release in pair bond formation and maintenance (*7, 8, 10*). Building on this, we used the subsecond temporal resolution of the fluorescent biosensor, GRAB_DA_, to query the specific dopaminergic dynamics that underlie partner and novel seeking and interaction in pair bonded prairie voles (*17*). Our within-animal design revealed enhanced dopamine release when voles experience cues predicting partner access and when they interact with their partner. In addition, consistent with other social operant reward tasks (18), we found that once the task is learned, dopamine is released not only during social access but also during events that predict social access (*2*). Partner-enhanced dopamine release was also diminished as a function of long-term partner separation. Together, these differences in partner- and novel vole-associated dopamine release may serve as a valuation signal to guide pair bond motivation and selective affiliative behavior.

One major experimental advantage of our approach is the ability to dissociate differences in dopaminergic signaling from differences in behavior. Pair bonds are hallmarked by striking differences in partner and novel-directed behavior; it can therefore be challenging to determine whether differences in behavior drive differential dopamine release or if differences in dopamine release drive differences in affiliative behavior. Our operant task effectively disentangles these two possibilities by querying dopamine release dynamics during equivalent behaviors and cues (e.g. lever press and door open) for the partner and novel vole. As voles exhibit the same amount of lever pressing during partner and novel trials, differences in dopamine release are also not attributable to differences in the amount of these behaviors. Our results indicate that differences in partner- and novel-associated dopamine release can appear independent of specific behaviors or their quantities. Real-time differences in dopamine release may thus encode key facets relevant to navigating social interactions, such as a prediction of the degree or type of social reward.

Enhanced partner-associated dopamine release and its erosion as a function of partner separation are consistent with a model of dopamine as a reward valuation signal. Specifically, enhanced dopamine release may help cement bonds by encoding highly rewarding aspects of partner interaction. Correspondingly, the blunting of dopamine responses following long term partner separation may reflect neuronal processes that enable the formation of a new bond by reducing the selective reward associations with a previously pair bonded partner. However, novel voles also elicit dopamine release and motivate similar amounts of lever pressing for access. Potential explanations for novel-elicited dopamine are likely complex. On explanation is that novel evoked dopamine release may encompass reward-related signaling tied to the potential for extra-pair copulations, which can increase fitness. Another possibility is that dopamine is released in response to a novel threat and a need to defend territories from intruders. The latter is consistent with prior work suggesting that dopamine signaling may mediate aggression and novelty detection in addition to social reward (*3, 7, 18, 20*). Ultimately, testing these models, which posit differential roles for partner- and novel-elicited dopamine release, will require functional manipulation.

While the present study provides novel insights into the real time dopamine dynamics that contribute to bond selectivity, examining aggregate changes in fluorescence as a proxy for dopamine release has some limitations. Specifically, we do not know whether there exists spatial or synapse-level specificity with respect to partner and novel-elicited dopamine release. Prior work suggests that the nucleus accumbens contains “hot spots” that mediate hedonic impact, motivation, reward and aversion (*21, 22*). A finer-grained analysis of dopamine dynamics may clarify whether bulk or localized differences in release are important for mediating differences in prosocial or territorial behavior, potentially through differential recruitment of dopamine D1- and D2-receptor-expressing medium spiny neurons. Another possibility is that dopamine-mediated effects on behavior are the result of different patterns of co-release of dopamine with glutamate or GABA or combined action with oxytocin or endogenous opioids, which represents another promising avenue of future research. Finally, while intriguing, additional work is needed to clarify how reductions in partner-elicited dopamine release following long-term separation may functionally contribute to loss adaptation in voles.

In sum, we have shown that dopamine release reflects the selective nature of pair bonds, with greater release associated with these highly rewarding relationships. This work has important implications for human relationships, suggesting that dopamine may confer selectivity by predicting and reinforcing the rewarding aspects of partner interaction and thereby cementing relationships over time. The erosion of release dynamics as a function of partner separation is consistent with a model in which pair bonds are assigned less motivational saliency following prolonged partner absence, thus providing a potential mechanism for overcoming loss. Together, this work suggests that real-time dopamine release dynamics are sensitive to experience and differentiate between different relationship types, acting to shape real-time social decision-making and behavior.

## Acknowledgements

We thank Yulong Li for providing the GRAB_DA_ plasmid. We thank the animal care staff at the University of Colorado Boulder. We also thank Kelly Winther, Katie Gallagher, and Kresil Gordon for managing the animal colony and providing experimental support. Additionally, Sarah Litz contributed to the scripts used to run the operant chambers. We took histology images on the Stem Cell Research and Technology Resource Center microscope. We thank Kelly Winther for slicing and mounting brain tissue and taking florescent microscopy images for histology. Finally, we thank the rest of the Donaldson lab for their feedback and support and the voles for their sacrifice. This work was supported by NIH awards R36MH129127 (to A. F. P.) & DP2MH119421 (to Z.R.D.).

## Author contributions

Z.R.D. and A.F.P developed experimental design. A.F.P. executed experiments. Z.R.D. and A.F.P. analyzed and interpreted data and wrote the manuscript. D.S.W.P, G.D.C, and R.T.C created the hardware and software for the operant chambers. D.S.W.P provided feedback on experimental design and edited the manuscript.

## Competing interest statement

Authors have no competing interest or disclosures.

## Materials and Methods

### Animals

Prairie voles were bred in house, originating from colonies housed at Cornell University, Emory University, and UC Davis, all of which originated from wild animals captured in Illinois. Animals were maintained at a temperature of 23-26°C on a 14:10 light:dark cycle. All procedures occurred during the light phase. Animals were given water and rabbit chow ad libitum (5326-3 by PMI Lab Diet). Rabbit chow was supplemented with sunflower seeds, dehydrated fruit bits, and alfalfa cubes. Home cages were enriched with cotton nestlets and plastic houses. At postnatal day 21, animals were weaned and placed into standard static rodent cages (17.5 l. x 9.0 w. x 6.0 h. in.) at a density of 2-4 same sex prairie voles. All females were sterilized via tubal ligation between the ages of postnatal day 72 and 96. At the onset of opposite-sex pairing, voles were placed into smaller static rodent cages (11.0 l. x 6.5 w. x 5.0 h. in.) where they remained until they were separated from their opposite-sex partner and placed in clean small rodent cages. All voles were between 94 and 118 days old at the onset of pairing. Procedures were approved by the University of Colorado Institutional Animal Care Use Committee.

### Surgical procedures

#### AAV infusion and ferrule implantation

Experimental animals underwent viral infusion and ferrule implantation surgery between 72 and 96 days of age. Voles were anesthetized with 1– 3% isoflurane at an oxygen flow rate of 1L/min in a head-fixed stereotactic frame (David Kopf, Tujunga, CA). Body temperature was maintained at 37°C using a closed loop heating pad with a rectal thermometer (David Kopf, Tujunga, CA). Eyes were lubricated with ophthalmic ointment (Puralube Vet Ointment). The fur was removed from the incision site using a shaver, and the wound area was disinfected with 70% ethanol and betadine. Briefly, the scalp and any connective tissue was removed above the frontal and parietal skull plates. Two 0.5 mm guide holes were drilled— each in the parietal plates —and anchoring screws were rotated into place. The head was leveled in the anterior-posterior plane, and a 0.5 mm hole was drilled at +1.6 mm AP and +1 mm ML. A Nanoject syringe (Drummond Scientific, Broomall, PA) was lowered and 200 nL of AAV vector was injected at a rate of 1nL/sec unilaterally at -5.0, -4.9, and -4.8mm DV for a total of 600 nL. The following vectors and titers were used: AAV1-hSyn-DA4.4 M205T(GRAB_DA_, plasmid gift from Dr. Yulong Li, packaged by Vigene): 1.015×1013 GC/ml; AAV2-hsyn-mCherry: 3×1012 GC/ml. The needle was left in place for 10 minutes following the last infusion. A fiber-optic ferrule (0.2 mm diameter, 5 mm long; Doric Lenses Inc) was slowly lowered into position until reaching a final placement of -4.8 mm DV. The ferrule and screws were affixed to the skull with Loctite 454 cured with dental cement liquid. The initial layer was covered with additional Loctite with black carbon powder (Sigma). Sustained Release Meloxicam (2 mg/kg), enrofloxacin (5 mg/kg), and saline (up to 3mL) were administered subcutaneously perioperatively for analgesia, to avoid bacterial infection at the wound site, and to prevent dehydration, respectively. Additionally, enrofloxacin (5 mg/kg), and saline (1mL) were administered subcutaneously for the three days following surgery. Animals were allowed to recover for at least 14 days prior to initiation of experiments. Ferrule placement and viral expression were confirmed posthumously (Fig 1B, C). Five animals were excluded due to placement outside of the NAc or misalignment of the ferrule with viral expression.

#### Tubal ligations

Females were tubally ligated to avoid confounds of pregnancy while keeping the ovaries hormonally intact. Tubal ligation was carried out under anesthesia during vector infusion and ferrule placement (described above). Briefly, hair was shaved at the incision site and the underlying skin was disinfected with betadine and 70% ethanol. A single incision was made in the midline of the back to provide access to the body cavity. The incision was pulled to one side until aligned above the ovary. A small incision was made into the body wall, the ovary was pulled through and bisected from the uterus via a cauterizer. The uterus and ovary were returned to the body cavity, the internal body wall was closed using an absorbable suture, the skin was pulled to the other side, and the procedure was repeated. The external incision was closed with staples that were removed 10-14 days later. Triple antibiotic ointment and lidocaine were placed on the closed wound.

### Behavioral Methods

#### Operant training and timeline

Custom operant chambers were as previously described in Brusman et al 2022 (15). Briefly, operant chambers contained 3 chambers separated by 2 motorized doors, one motorized pellet dispenser and trough, and 3 separate retractable levers (one for each type of reward) (Fig 1D). Chambers were constructed from a mix of laser cut acrylic and 3D printed ABS plastic. A bill of materials and chamber designs can be accessed at https://github.com/donaldsonlab/Operant-Cage/tree/main/V2.

The apparatus was controlled via custom scripts and code (https://github.com/dprotter/RPi_Operant) run on Raspberry Pi computers (Raspberry Pi Foundation). Servos were controlled via an Adafruit HAT (Adafruit 2327). Each apparatus was controlled by a corresponding Raspberry Pi. Food rewards were 20 mg pellets (Dustless Precision Pellets Rodent Grain-Based Diet; VWR 89067-546) delivered to a trough. Pellet dispensing and retrieval was detected by an IR beam break in the trough. Tones were generated via PWM on the Raspberry Pi (pigpio), and played through an amplified speaker (Adafruit 3885).

Voles were paired more than 14 days prior to the onset of operant training, sufficient time for stable pair bonding to occur (15, 23). Operant training and testing consisted of 6 days of magazine training, 8 days of operant food delivery, and 6 days of operant social reward delivery (Fig. S1).

#### Food Magazine Training

Animals underwent 15 trials per day, the goal of which was to learn associations between the lever, tone, and food reward. For each trial, a tone was played to indicate the start of the trial (5,000 Hz, 1s). The food lever was then extended for 2 seconds, a pellet cue (2,500 Hz, 1s) was played, and a pellet was delivered to the trough. The lever was retracted 2 seconds later. If an animal pressed within the first 2 seconds of lever access, a pellet was immediately delivered. No more than 1 pellet was delivered per trial. Total trial time was 90s.

#### Operant food delivery

Animals underwent 20 trials per day for 8 days (Fig S1B). During the first two days (training), pellet delivery was not contingent on lever pressing. During each trial, a tone was played to indicate the start of the trial (5,000 Hz, 1s). The food lever was then extended for 30 seconds. After 30 seconds, the lever was retracted if the vole did not press the lever, a pellet cue (2,500 Hz, 1s) was played, and a pellet was delivered to the trough. If the vole pressed the lever within 30 seconds, a pellet cue (2,500 Hz, 1s) was played, and a pellet was immediately delivered to the trough. Thus, a single pellet was dispensed on every trial after 30s of lever presentation, but lever pressing elicited an immediate pellet dispense. During days 3-8 of training, pellet delivery was contingent on lever pressing. The lever was extended for a maximum duration of 120s. During each trial, a tone was played to indicate the start of the trial (5,000 Hz, 1s). After 120 seconds, the lever was retracted if the vole did not press the lever and no pellet was dispensed. If the vole pressed the lever within 120 seconds, a pellet cue (2,500 Hz, 1s) was played, and a pellet was delivered to a trough. In order to provide a window to observe anticipatory behavior and dopamine release, animals experienced a delay between lever pressing and food reward as follows: days 1-5: no delay, days 6-8: 5 second delay. The intertrial interval for all trials was 45 sec.

### Enhanced dopamine during partner seeking

#### Operant social access

Animals underwent 20 trials of social training per day, the design of which mirrored days 3 - 8 of operant food delivery except the lever press resulted in social access to the partner or a novel, opposite-sex vole. Experimental voles were administered alternating sets of 5 trials for each door, starting with the partner door (i.e. 5x partner, 5x novel, 5x partner, 5x novel) (Fig 1E, F). The partner and novel stimulus animals were tethered at opposite ends of the apparatus and farthest from the doors in a similar fashion to the partner preference test (below). The tethering location of partner and novel vole remained consistent across days. To avoid a potential unintended bias in lever pressing, we assign the lever farthest from the door to provide access to the partner or novel, respectively (Fig 1D). A new novel vole was used each day of operant social access and during the probe trial after separation. On all training days, a tone was played to indicate the start of the trial (5,000 Hz, 1s). After 120 seconds, the lever was retracted. If the lever was pressed within 120 seconds, social access was granted, and at the end of the trial, a door close tone was played (7,000 Hz, 1s) and the door was closed. If needed, subjects were manually returned to the center chamber immediately after the chamber closed. The duration of social interaction received was dependent on how quickly the vole pressed the lever. Each trial was a maximum of 120 seconds, and the amount of social interaction was 120 second minus the latency to press the lever. All trials had an intertrial interval of 45 seconds. The door was opened without any delay after lever press on days 1 - 3 and following a 5 second delay on days 4 - 6.

#### Fiber photometry and data analysis

Subjects were habituated to the patch cable for 6 days in an open field chamber prior to the onset of operant training/testing. Subjects were briefly anesthetized (<30s) to attach patch cables prior to recording and allowed 10 minutes to recover prior to operant testing.

Fluorescence was acquired using the Neurophotometrics (NPM) V2 system with 200uM core optical fibers purchased from Doric Lenses. Data was acquired using Bonsai During photometry recordings, light was delivered alternating between 470 nm, 560 nm, and 415 nm at a framerate of 180 frames per second. The LED power for each wavelength was set to 50uW at the optical fiber tip to reduce photobleaching. Signals were analyzed using a MATLAB script. To correct for photobleaching and motion artifacts, the 560 nm signal was fit to the 470 nm, then this fit was subtracted from the 470 nm signal. Z-Scores were calculated as (fitted signal - baseline)/(baseline standard deviation) where the baseline for all events and behaviors was -8 to -3 seconds prior to lever extension. The area under the curve was calculated as the average Z-Score values during the time period of interest (typically 2 sec after the event).

We time-locked operant events (extension of levers, lever pressing, chamber opening, chamber entry, pellet dispense, pellet retrieval) to the fluorescence signal using two microcontrollers. Bonsai cannot run on the Raspberry Pi 3 B+ operating system that is used to control the operant chamber hardware, so we directed a Raspberry Pi to send a serial signal to an Arduino Uno microcontroller, which has communication functionality with Bonsai.

All behaviors that occurred after crossing into the chamber were hand scored using BORIS, a behavior event scoring software (24). Behaviors performed by the subject directed towards the partner or novel on days 4-6 of social operant access were hand scored. We examined the following behaviors: non-contact investigation, head sniffing, body sniffing, anogenital sniffing, huddling, allogrooming, defensive posture, and attacking.

#### Partner preference test

Partner preference tests were performed as described in Scribner et al. 2020 (23). Briefly, both partner and novel animals were tethered to the end walls of three-chamber plexiglass arenas (76.0 cm long, 20.0 cm wide, and 30.0 cm tall). Tethers consisted of an eye bolt attached to a chain of fishing swivels that slid into the arena wall. Animals were briefly anesthetized with isoflurane and attached to the tether using a zip tie around the animal’s neck. Two pellets of rabbit chow were given to each tethered animal and water bottles were secured to the wall within their access while tethered. After tethering the partner and novel animals, experimental animals were placed in the center chamber of the arena. At the start of the test, the opaque dividers between the chambers were removed, allowing the subject to move freely about the arena for three hours. Overhead cameras (Panasonic WVCP304) were used to video record eight tests simultaneously.

The movement of all three animals in each test was scored using TopScan High-Throughput software v3.0 (Cleversys Inc) using the parameters from Ahern et a., 2009 (13). Behavior was analyzed using a Python script developed in-house (https://github.com/donaldsonlab/Cleversys_scripts) to calculate the time spent huddling with the partner or novel. The partner preference score was calculated as (partner huddle time/[partner huddle time + novel huddle time]) × 100%.

### Brain collection

Upon completion of all experimental sessions, voles were transcardially perfused with 4% paraformaldehyde in phosphate buffered saline. The head was removed and post-fixed for 24 hours in 4% paraformaldehyde before extracting the brain. The brain was equilibrated in 30% sucrose, sectioned in 50 μm slices using a sliding freezing microtome (Leica), and mounted on slides. Ferrule placement was drawn onto corresponding mouse atlas sections.

### Statistical Analysis

All statistical analyses were carried out using Graphpad PRISM 9.3.1 and SPSS 29.0.0.0. Events corresponding to learning the operant task (number of lever presses, latency to lever press, latency to cross into a social chamber, and pellet retrieval were analyzed using a one-way repeated measures ANOVA or mixed model ANOVA to assess effect of day (Fig 1, Fig S1) or two-way repeated measures ANOVA (Fig 2) to examine main effects of day and partner versus novel. Z-scored fluorescence area under the curve comparisons for day 1 and day 6 (Fig 1) or partner and novel (Fig 2, 3, 4) were analyzed using a paired or unpaired t-test (latter only Fig 3H). Differences in Z-scored fluorescence area under the curve elicited by food and social operant events were analyzed using a one-way repeated measures ANOVA with a Tukey’s post-hoc test (Fig S1). Partner preference was analyzed using a one sample t test relative to an expected null 50 percent preference (no preference) (Fig 2B). Differences in durations of behaviors demonstrated towards partners and novels were analyzed using a paired or unpaired t-test (latter only for Fig 3E). Finally, cumulative bouts of behaviors demonstrated towards partners and novels were analyzed using a Log-rank (Mantel-Cox) test. See Supplement table 1 for a full list of statistics information.

**Figure S1:**
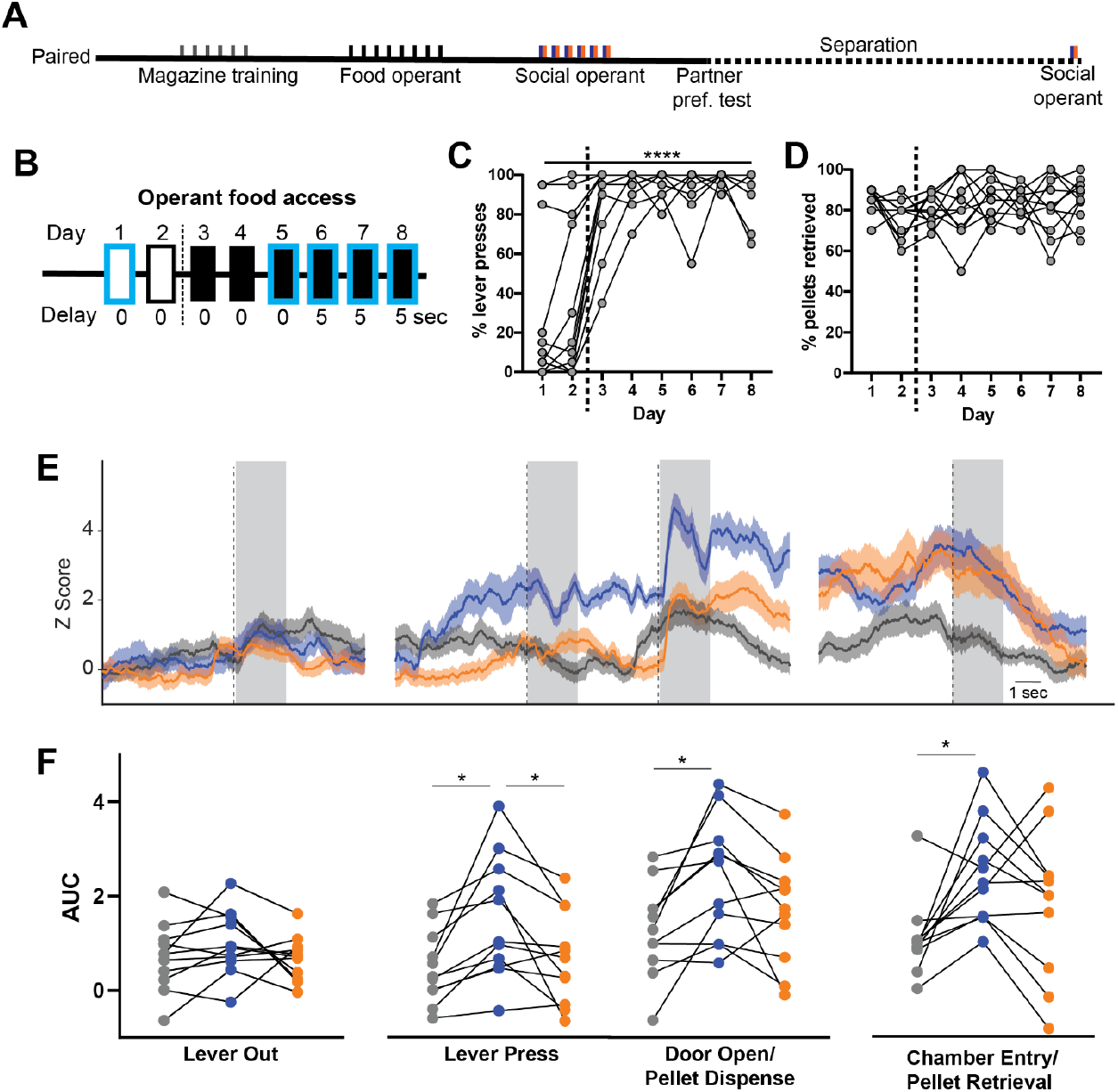
Enhanced dopamine dynamics for partner operant events compared to food. A) Timeline for experiment. B) Timeline for operant food access. Blue outline indicates fiber photometry recording of dopamine release. Pellet delivery was contingent on lever pressing from Day 3 onwards (black filled rectangles). C) Lever pressing increased across days, particularly after lever contingency was implemented on day 3 (one-way repeated-measures ANOVA p <0.0001) D) Pellet retrieval remained high across all training days independent of lever pressing contingency (oneway repeated-measures ANOVA p = 0.4034). E) Representative GRAB_DA_ fluorescence in response to the onset of events (dotted line). The fluorescence is Z-scored relative to baseline. F) Area under the curve (AUC) for 2 sec after event onset (shaded regions in E). For all operant events aside from lever extension (lever out), partner-associated events elicit greater dopamine release than the same events directed towards food or the novel. One-way repeated-measures ANOVA comparing partner, novel, and food. (Lever out p = 0.3122. Lever press p = 0.0079. Posthoc Tukeys test comparing partner and food p = 0.0135; novel and food p = 0.9938; partner and novel p = 0.0461. Door open/pellet dispense p = 0.0304. Posthoc Tukeys test comparing partner and food p = 0.0174; novel and food p = 0.6368. Chamber entry/pellet retrieval p = 0.0091. Posthoc Tukeys test comparing partner and food p = 0.0166; novel and food p = 0.1689.) * = p < 0.05; ** = p < 0.01; *** = p < 0.001.

## Notes

### Competing Interest Statement

The authors have declared no competing interest.

