## Supplementary material for "Nucleus accumbens dopamine release reflects the selective nature of pair bonds": Table S1

| Measurement | Statistical Test | Comparison | ° of freedom, error | F or T | p | * | Effect size (cohen's d / eta squared) | Fig. | Post hoc tests |
|---|---|---|---|---|---|---|---|---|---|
| Avg number of lever presses (partner and novel aggregated) | One-way repeated-measures ANOVA | days | 5 | 6.139 | 0.0047 | ** | 0.260303 | 1G | |
| Avg latency to press (partner and novel aggregated) | Mixed effects analysis | days | 5 | 1.9 | 0.1783 | ns | 0.165981 | 1H | |
| Avg latency to cross after chamber open (partner and novel aggregated) | Mixed effects analysis | days | 5 | 8.217 | 0.0018 | ** | 0.419672 | 1I | |
| AUC Lever out | Paired t-test | Day 1 and Day 6 | 10 | 2.716 | 0.0217 | * | 1.189837 | 1K | |
| AUC lever press | Paired t-test | Day 1 and Day 6 | 10 | 2.948 | 0.0146 | * | 1.240892 | 1K | |
| AUC chamber open | Paired t-test | Day 1 and Day 6 | 10 | 4.696 | 0.0008 | *** | 1.757846 | 1K | |
| AUC chamber cross | Paired t-test | Day 1 and Day 6 | 10 | 1.474 | 0.1711 | ns | 0.521262 | 1K | |
| Partner Preference | One sample t relative to 50% | % huddle compared to 50% (null) | 10 | 2.895 | 0.016 | * | 0.9641 | 2B | |
| Avg number of lever presses | Two-way RM ANOVA | partner vs novel | 1,10 | 1.592 | 0.2357 | ns | 0.00893 | 2C | |
| | | days | 5, 50 | 6.139 | 0.0002 | *** | 0.19339 | | |
| | | day X partner/novel | 5,50 | 6.273 | 0.0001 | *** | 0.092736 | | |
| Avg latency to lever press | Mixed effects analysis | partner vs novel | 1,10 | 1.683 | 0.156 | ns | 0.020637 | 2D | |
| | | days | 5, 50 | 0.8228 | 0.2185 | ns | 0.065997 | | |
| | | day X partner/novel | 5,45 | 0.2377 | 0.9438 | ns | 0.006766 | | |
| Avg latency to cross after chamber open | Mixed effects analysis | partner vs novel | 1,10 | 0.5406 | 0.4791 | ns | 0.001608 | 2E | |
| | | days | 5, 50 | 10.4 | <0.0001 | **** | 0.347764 | | |
| | | day X partner/novel | 5,45 | 0.2369 | 0.9441 | ns | 0.006754 | | |
| AUC Lever out | Paired t-test | Partner and Novel | 10 | 1.382 | 0.1972 | ns | 0.549289 | 2G | |
| AUC lever press | Paired t-test | Partner and Novel | 10 | 2.791 | 0.0191 | * | 0.718155 | 2G | |
| AUC chamber open | Paired t-test | Partner and Novel | 10 | 2.307 | 0.0438 | * | 0.572761 | 2G | |
| AUC chamber cross | Paired t-test | Partner and Novel | 10 | 1.515 | 0.1606 | ns | 0.457808 | 2G | |
| AUC chamber cross sec 4-5 | Paired t-test | Partner and Novel | 10 | 3.26 | 0.0086 | ** | 0.792312 | 2G | |
| Body sniff duration | Paired t-test | Partner and Novel | 5 | 3.549 | 0.0164 | * | 1.338562 | 3A | |
| Body sniff cumulative bout # | Log-rank (Mantel-Cox) test | Partner and Novel | 1 | 17.67 | <0.0001 | **** | na | 3B | |
| Body sniff AUC | Paired t-test | Partner and Novel | 5 | 8.974 | 0.0003 | *** | 0.950428 | 3D | |
| Huddle duration | Unpaired t-test | Partner and Novel | 7 | 2.423 | 0.0459 | * | 2.047358 | 3E | |
| Huddle cumulative bout # | Log-rank (Mantel-Cox) test | Partner and Novel | 1 | 16.25 | <0.0001 | **** | na | 3F | |
| Huddle AUC | Unpaired t-test | Partner and Novel | 7 | 3.268 | 0.0137 | * | 2.269937 | 3H | |
| Non-contact investigation duration | Paired t-test | Partner and Novel | 10 | 2.795 | 0.019 | * | 1.004982 | 3I | |
| Non-contact investigation cumulative bout # | Log-rank (Mantel-Cox) test | Partner and Novel | 1 | 6.565 | 0.0104 | * | na | 3J | |
| Non-contact investigation AUC | Paired t-test | Partner and Novel | 10 | 1.085 | 0.3033 | ns | 0.460414 | 3L | |
| Partner lever out AUC | Paired t-test | Pre and post separation | 10 | 2.88 | 0.0164 | * | 0.880079 | 4B | |
| Novel lever out AUC | Paired t-test | Pre and post separation | 10 | 0.717 | 0.4898 | ns | 0.303251 | 4B | |
| Partner lever press AUC | Paired t-test | Pre and post separation | 9 | 0.0428 | 0.9668 | ns | 0.020193 | 4C | |
| Novel lever press AUC | Paired t-test | Pre and post separation | 7 | 0.9486 | 0.3744 | ns | 0.307337 | 4C | |
| Partner door open AUC | Paired t-test | Pre and post separation | 9 | 2.403 | 0.0397 | * | 1.091031 | 4D | |
| Novel door open AUC | Paired t-test | Pre and post separation | 7 | 0.5486 | 0.6003 | ns | 0.099697 | 4D | |
| Partner chamber entry AUC | Paired t-test | Pre and post separation | 9 | 2.705 | 0.0242 | * | 0.975496 | 4E | |
| Novel chamber entry AUC | Paired t-test | Pre and post separation | 7 | 1.479 | 0.1826 | ns | 0.316318 | 4E | |

| | | | | | | | | | |
|---|---|---|---|---|---|---|---|---|---|
| Post-separation lever out AUC | Paired t-test | Partner and novel | 10 | 0.0379 | 0.9705 | ns | 0.014214 | 4F | |
| Post-separation lever press AUC | Paired t-test | Partner and novel | 7 | 1.502 | 0.1769 | ns | 0.605873 | 4F | |
| Post-separation door open AUC | Paired t-test | Partner and novel | 7 | 0.639 | 0.5432 | ns | 0.272457 | 4F | |
| Post-separation chamber entry AUC | Paired t-test | Partner and novel | 7 | 2.039 | 0.0808 | ns | 0.787113 | 4F | |
| % lever presses for food pellet | One-way repeated-measures ANOVA | Days | 7 | 23.09 | <0.0001 | **** | 0.697781 | S1C | |
| % pellet retreivals | One-way repeated-measures ANOVA | Days | 7 | 1.03 | 0.4034 | ns | 0.093414 | S1D | |
| Lever out AUC | One-way repeated-measures ANOVA | Partner, novel and food | 2, 10 | 1.224 | 0.3122 | ns | 0.109034 | S1F | Tukeys: df = 10; partner vs food: p = 0.276; novel vs food: p = 0.9938. |
| Lever press AUC | One-way repeated-measures ANOVA | Partner, novel and food | 2, 10 | 6.338 | 0.0079 | ** | 0.387943 | S1F | Tukeys: df = 10; partner vs food: p = 0.0135; novel vs food: p = 0.8184; partner vs novel: p = 0.0461. |
| Door open/pellet dispense AUC | One-way repeated-measures ANOVA | Partner, novel and food | 2, 10 | 4.75 | 0.0304 | * | 0.322031 | S1F | Tukeys: df = 10; partner vs food: p = 0.0174; novel vs food: p = 0.6368 |
| Chamber cross/pellet retreive AUC | One-way repeated-measures ANOVA | Partner, novel and food | 2, 10 | 6.005 | 0.0091 | ** | 0.375195 | S1F | Tukeys: df = 10; partner vs food: p = 0.0166; novel vs food: p = 0.1689 |
